# Assessing robustness against potential publication bias in coordinate based fMRI meta-analyses using the Fail-Safe N

**DOI:** 10.1101/189001

**Authors:** Freya Acar, Ruth Seurinck, Simon B. Eickhoff, Beatrijs Moerkerke

**Affiliations:** Faculty of Psychology and Educational Sciences, Ghent University, Belgium; Institute of Systems Neuroscience, Medical Faculty, Heinrich Heine University Düsseldorf, Düsseldorf, Germany, Institute of Neuroscience and Medicine, Brain & Behaviour (INM-7), Research Centre Jülich, Jülich, Germany

**Author notes:** Corresponding author: Freya Acar, Henri Dunantlaan 2, Ghent, Belgium, telefacsimile: 00329 264 64 87.

**Keywords:** publication bias, coordinate based meta-analysis, fMRI, file-drawer, robustness, ALE, PET

## Abstract

The importance of integrating research findings is incontrovertible and coordinate based meta-analyses have become a popular approach to combine results of fMRI studies when only peaks of activation are reported. Similar to classical meta-analyses, coordinate based meta-analyses may be subject to different forms of publication bias which impacts results and possibly invalidates findings. We develop a tool that assesses the robustness to potential publication bias on cluster level. We investigate the possible influence of the file-drawer effect, where studies that do not report certain results fail to get published, by determining the number of noise studies that can be added to an existing fMRI meta-analysis before the results are no longer statistically significant. In this paper we illustrate this tool through an example and test the effect of several parameters through extensive simulations. We provide an algorithm for which code is freely available to generate noise studies and enables users to determine the robustness of meta-analytical results.

## Main text

### 1. Introduction

Functional magnetic resonance imaging (fMRI) continues to contribute greatly to the knowledge about the location of cognitive functions in the brain (Logothetis, 2008). Neuroimaging studies are valuable sources of information for functional organisation of the brain but often face substantial challenges. A huge variety in employed experimental conditions and analytical methods exist, complicating finding consistency across studies and paradigms. Several different implementations and task contrasts might be applied while exploring a certain paradigm, using different analysis toolboxes, pipelines and statistical thresholds. Furthermore, fMRI studies are relatively expensive which often limits the size of studies (Carp, 2012) and small sample sizes lead to low statistical power to detect true activation (Durnez et al., 2014). Further progress in understanding human brain function will therefore also require integration of data across studies using meta-analyses, which can increase both power and reproducibility (Sutton et al., 2000).

fMRI studies capture information about more than 100.000 voxels in the brain, however, the most prevailing trend to report results remains providing coordinates of statistically significant local maxima, peaks or foci (Maumet et al., 2016). Several coordinate-based meta-analysis techniques have been developed to combine results of studies that employ this censored reporting of results. First Activation Likelihood Estimation (ALE; Eickhoff et al., 2009; Turkeltaub et al., 2012; Eickhoff et al., 2012) was developed, which is used most (see Figure 1). Later on, Seed-based d Mapping (Radua and Mataix-Cols, 2012) and multilevel kernel density analysis (MKDA; Wager et al., 2009) arose. As MKDA, ALE uses as input xyz-coordinates of the peaks (foci) reported by the individual studies and determines at which location the convergence of activation is larger than can be expected by chance. Seed-based d mapping also takes peak height into account and offers the possibility of entering entire t-maps into the meta-analysis. An advantage to ALE is that it is accompanied by the BrainMap database (Laird et al., 2005; Fox and Lancaster, 2002; Fox et al., 2005). The BrainMap database is a collection of over 3000 papers, containing information about e.g. experimental conditions, subjects and allows to extract peak locations in the format adapted to the ALE algorithm.

**Figure 1.**
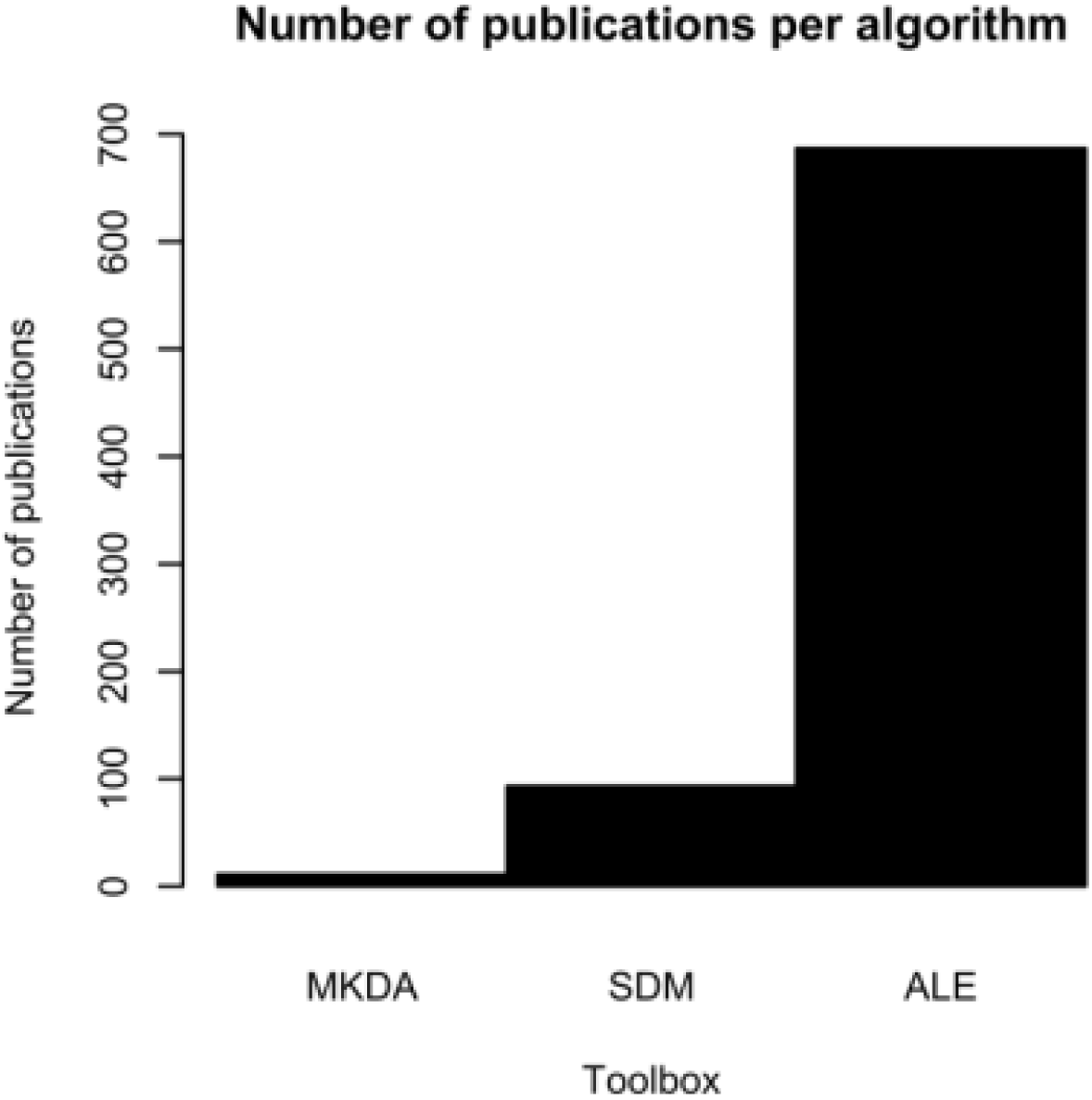
Number of publications per algorithm found after a web of science and web search executed on 21 April 2017. On web of science search terms were GingerALE, ALE meta-analysis for the ALE-algorithm, MKDA and MKDA meta-analysis for the MKDA algorithm and SDM meta-analysis, ES-SDM meta-analysis and seed-based d mapping for the seed-based d-mapping algorithm. The websites of the algorithms were also searched for corresponding publications (http://brainmap.org/ for the ALE-algorithm and http://www.sdmproject.com/ for seed-based d mapping).

The ALE algorithm offers a nifty workaround for the lack of data reported in an fMRI study. It models spatial uncertainty by constructing Gaussian kernels around the foci, with the size of the kernels depending on the individual study sample size. In large studies, a higher statistical power is attained and therefore there is smaller spatial uncertainty. This results in smaller kernels with higher values per voxel. By doing so it partially addresses the problem of bias that occurs due to the lack of reported information on activation, i.e. within-study publication bias. However, in general, meta-analyses are also vulnerable for between-study publication bias. This occurs when studies have a smaller chance of getting published if they fail to show certain results, this is commonly known as the file drawer problem (Rothstein et al., 2006). The file-drawer problem can affect the outcome of a meta-analysis by overestimating effects, as no contra-evidence is taken into account. Up to now, methods assessing the presence of such publication bias for coordinate-based meta-analysis techniques are scarce. As seed-based d mapping takes peak height into account it offers the opportunity for constructing a funnel plot, but for the ALE algorithm and MKDA no tools are available.

The file-drawer problem in classical meta-analyses is known for quite some time and has been extensively discussed in literature (Rothstein et al., 2006; Rosenthal, 1979; Kicinski, 2014). Rosenthal (1979) proposed the Fail-Safe N to assess to what extent publication bias may influence the results of a meta-analysis. It quantifies the number of null studies (i.e. studies without statistically significant results) that are needed to alter a meta-analytical result from statistically significant to statistically nonsignificant. The Fail-Safe N is based on the size of the effect resulting from the meta-analysis, the number of studies entered into the meta-analysis and a statistical threshold. If a meta-analysis consists of studies with large sample sizes and results in a large effect, the associated Fail-Safe N will also be large, indicating that it would take a large amount of unpublished null studies to alter the conclusions of the meta-analysis. Apart from being a measure for publication bias the Fail-Safe N also functions as measure of robustness as it quantifies the number of studies that are needed to alter results, with a low number being indicative for low robustness. In that sense, it can be used as an indication for how robust results are against publication bias. If the expected amount or percentage of null studies that remain in the file drawer is known, it is possible to determine whether results remain valid if these null studies would be entered into the meta-analysis. The use of Fail-Safe N in the context of publication bias for meta-analyses has been heavily debated (Rothstein et al., 2006; Hsu, 2002; Scargle, 2000; Schonemann and Scargle, 2008). One of the main critiques is the variety of methods that exist to compute the Fail-Safe N and the fact that different definitions for null studies circulate (Rosenthal, 1979; Orwin, 1983; Hsu, 2002). Depending on the choice of method and the definition of a null study the results of the Fail-Safe N can vary greatly. Furthermore, the large emphasis on achieving statistical significance instead of focusing on effect sizes has been pointed out (Rothstein et al., 2006). Orwin (1983) subsequently adapted the Fail-Safe N to measure to what extent effect sizes are altered by adding null studies in a meta-analysis. Despite these criticisms, we argue that such measure is useful to assess robustness in coordinate based meta-analyses of fMRI studies given the limited amount of information that is available for each individual study (i.e. the number of subjects and the location of the peaks or foci). In this paper, we propose a method for computing the Fail-Safe N for ALE meta-analyses but the concept is also more generally applicable for use in other coordinate based meta-analyses. The general idea is to add studies that do not report activation in a target area to an existing meta-analysis to determine the amount of contra-evidence that is needed before a statistically significant cluster is no longer deemed active. We will call the studies with contra-evidence “noise studies”. These studies contain activation but this activation is randomly distributed throughout the brain. This is explained in more detail in section 2.2, adding noise studies to an ALE meta-analysis. The results of this test can be interpreted in two ways. On the one hand, one wants to be able to add a minimal amount of noise studies to ensure that the results are robust against missing information, e.g. because of publication bias. On the other hand, if a large amount of noise studies can be added with only a small percentage of studies contributing to a statistically significant target cluster this implies that results are driven by a small proportion of all studies.

The recent mass simulation paper of Eickhoff et al. (2016) provides end users with clear and motivated guidelines for performing a meta-analysis with the ALE algorithm (Eickhoff et al., 2009; Turkeltaub et al., 2012; Eickhoff et al., 2012), recommending cluster-level family-wise error correction for moderate effects and a set size of at least 20 studies. With the use of Fail-Safe N, we aim to complement these general guidelines by providing an added value to the interpretation of brain regions selected as statistically significant by the ALE algorithm. This is achieved by taking into account specific characteristics of the studies in the meta-analysis set in the construction of the set of noise studies used to calculate the Fail-Safe N.

We introduce and illustrate the method to calculate the Fail-Safe N with an example meta-analysis and perform extensive simulations to test the influence of number of peaks, sample size and thresholding method on the outcome of the Fail-Safe N. Based on the methods outlined in the paper, we also provide a script that allows a researcher to generate noise studies based on the specific properties of their original meta-analysis, namely sample sizes and number of peaks of the individual studies. Researchers can use these noise studies to compute a Fail-Safe N adapted to their specific meta-analysis. Doing this provides a greater insight in the obtained results and interpretation.

### 2. Materials and methods

The Fail-Safe N (FSN) has been introduced in classical meta-analysis literature to quantify the robustness of a meta-analysis. It assumes that non-significant studies are still in the “file drawer” and assesses the required number of studies to change the conclusions of the meta-analysis (Rosenthal, 1979). First, we will briefly discuss the original concept and interpretation of the FSN in a classical meta-analysis. Next, we will discuss how to obtain the FSN for an ALE meta-analysis. We will then show the process of computing the FSN through an example of a real meta-analysis and through a set of extensive simulations.

#### 2.1 The Fail-Safe N

Regardless of the area of research, studies with statistically significant results are more likely to get published than studies with non-significant results (Dickersin et al., 1987). This implies that the published literature is not an accurate representation of executed research and results of meta-analyses may be biased. Faced with this problem Rosenthal developed a method to determine the amount of studies remaining in the file drawer necessary for an effect to become statistically non-significant. Even though he did not propose a statistical threshold to decide whether the FSN is big enough or not, he suggested that a FSN smaller than 5k + 10 (with k the amount of studies in the meta-analysis) should raise concern. If only studies that show statistically significant results get published and no true effect is present, then it is possible that all studies in the meta-analysis are false positives. Based on the assumed severity of the file-drawer problem and the publishing rate, every researcher should determine a minimum FSN adapted to the area of research. Assume *k* independent studies where for each individual study *i, a z*_*i*_,-value and one-tailed p-value *p*_*i*_ is used to test the null hypothesis *H*_*0*_: *ϑ*_*i*_ = 0, with *ϑ* representing an effect size. The original calculation of the FSN is based on computing an overall p by adding Z’s (Cochran, 1954):

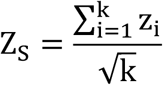

Based on this formula, the number of studies N with Z = 0 that are necessary for ZS to become smaller than the z-value cutoff Zα can be attained.

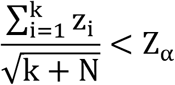

This eventually leads to the formula for the FSN

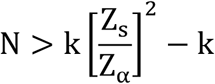

We now illustrate the FSN with an example from the book “Publication Bias in Meta-Analysis” (Rothstein et al., 2006). The analysis is based on a dataset from Raudenbush (1984) on teacher expectancy, a paradigm where the influence of teacher expectancy on intelligence test performance is measured. The dataset consists of 19 randomized studies, with one-tailed probability values ranging from 0.0158 to 0.9255. The Stouffer sum of 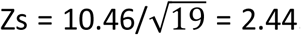. If we use this information to compute the Fail-Safe N we get:

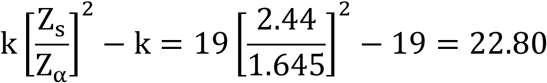

This means that 23 studies with a z-value of 0 need to be added to render the previously statistically significant results non-significant. This is less than the proposed 105 and caution is advised when interpreting the results about the teacher expectancy effect. Imagine if a little bit over 50% of the executed studies remain in the file-drawer because they showed no statistical significant effect. Adding these studies to the metaanalysis would reduce the sum of Zs to 0. In another example from the original paper of Rosenthal (1979) on interpersonal self-fulfilling prophecies the FSN is N = 3263 where k = 94. We can be more confident about these results as it seems highly unlikely that 3263 studies remain unpublished.

In classic meta-analysis the FSN is calculated by computing the average z-value or effect size and deducting the amount of studies that can be added before this summary statistic is below a certain threshold. In the meta-analysis of fMRI studies however we want to compute the FSN for a cluster of activation. Every cluster consists of several voxels, each with their own statistical value. Combining these would be challenging. Furthermore typically only the location (and sometimes the height) of voxels that exceed a statistical threshold are reported, which makes computing a summary value even more challenging. The ALE algorithm solves this by eventually computing a union of activation instead of a sum or average. Therefore the classic implementation of the FSN cannot be applied here. In the next section we will clarify how the FSN for a meta-analysis of fMRI studies is defined and computed.

#### 2.2 Adding noise studies to an ALE meta-analysis

As a meta-analytic method for coordinate based meta analyses, the ALE algorithm determines whether the spatial convergence of foci across studies is larger than can be expected by chance. An overview of the method is illustrated in Figure 2. While the result of an fMRI data analysis is a statistical parametric map, often only peak locations are reported. In the ALE algorithm these reported foci are entered into an empty brain by assigning them a value of 1, all other voxels have a value of 0. For every study a map is constructed with its reported foci. In a next step this value of 1 is smeared out by smoothing to neighbouring voxels by means of a Gaussian kernel of which the size depends on the sample size of the study. Small studies have less statistical power and more spatial uncertainty and therefore kernel size increases with decreasing sample size. The resulting map is called a modelled activation (MA) map and represents the probability of true activation being present in that voxel. Subsequently the MA-maps are combined into one ALE-map by taking the union of probabilities (equation 1). For every voxel with coordinates x, y, z, the ALE value in the map is the result of taking the union of the MA-values of that voxel over the k studies entered into the meta-analysis,

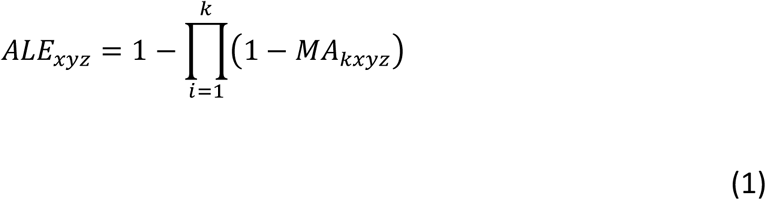

Thresholding of the resulting ALE map can be done both voxelwise (uncorrected, or with multiple testing corrections to control the False Discovery Rate FDR or the Family Wise Error FWE rate) or on cluster-level. In the case of voxel-level thresholding, a null distribution is obtained by constructing a histogram with all possible ALE values. This is done by computing all possible combinations from the values in the MA-maps. This histogram is used to calculate the uncorrected, FDR or FWE-corrected threshold. On top of a voxel-level threshold, a cluster-level threshold can be added by randomly distributing the MA-values throughout the brain, performing the same voxel-level threshold and listing the size of the largest resulting cluster. This process is repeated (e.g. 1000 times) and the distribution of cluster sizes is used to compute a cluster-level threshold. The authors of the ALE algorithm advise to employ a voxel-or cluster-level family-wise error correction (FWE; Eickhoff et al., 2016).

The results of an ALE meta-analysis consist of clusters where all voxels survived a statistical threshold.

**Figure 2.**
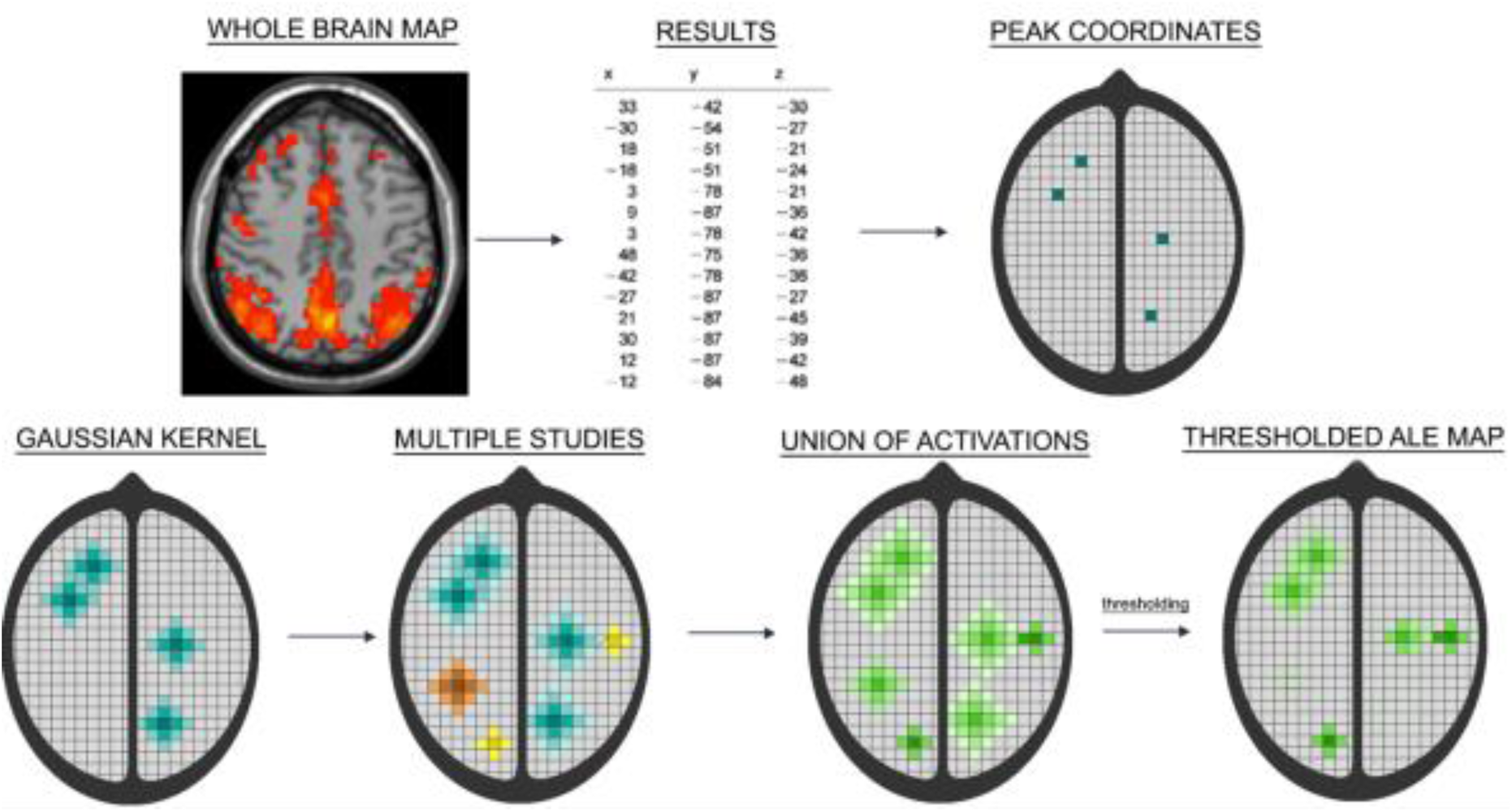
Step-by-step overview of the ALE algorithm. From an entire Statistical Parametric Map (SPM) usually only peak coordinates (foci) are reported. These are entered into an empty brain for every study individually. Spatial uncertainty is accounted for by constructing Gaussian kernels around the foci of which the size depends on the sample size of that study (smaller studies have more spatial uncertainty and larger kernels). The result of this process is a set of MA-maps. An ALE-map is constructed by calculating the union of the MA-maps. Eventually the ALE-map is thresholded to determine at which locations the convergence of activation is larger than can be expected by chance.

Cluster locations, size and anatomical region can be viewed and the number of contributing peaks per study can be computed. This allows to verify how the activated cluster came about. Knowing the number of studies and the number of peaks per study that contributed to a cluster can be very informative. Large clusters can either originate from a large number of studies or from a small number of studies with each a large number of peaks. A cluster from the first scenario, with a lot of contributing studies, seems more relevant and more reliable than a cluster with only a few contributing studies. Adding information about the number of participants in the contributing studies aids the interpretation of a cluster even more, with large studies providing more evidence for true activation, due to higher power, than small studies.

To determine the amount of noise studies that can be added to an existing meta-analysis without altering results for a given region with statistically significant activation, studies that show no effect in that region need to be simulated. In the ALE algorithm it is not possible to enter studies without any peaks as null studies because adding studies without foci does not change the results of the meta-analysis. Referring to equation (1), if an MA__kxyz__ of 0 is entered into this formula, the second part is merely multiplied by 1. Therefore these noise studies are conceptualized as studies with activation in a random location. Eickhoff and colleages (2016) have shown that adding random coordinates to an ALE meta-analysis barely changes the resulting ALE values. The threshold however does change, with larger ALE values necessary for statistical significance.

An R programme that can be used to generate noise studies can be found on Github (https://github.com/NeuroStat/GenerateNull). Researchers read in the same list of foci that was entered into ALE. From this list the number of peaks and participants per study are saved, each in an individual vector. The researcher has the possibility to indicate the number of noise studies he or she wants to generate. If no number is entered 10k (with k the number of experiments in the original meta-analysis) noise studies are generated. The number of peaks and number of participants of these noise studies are randomly sampled from the previously saved vectors. If the original meta-analysis only contains studies with less than 20 subjects, the noise studies will also have less than 20 subjects. Likewise, if the studies in the meta-analysis reported between 10 and 20 foci, the simulated noise studies will also report between 10 and 20 foci. The location of the peaks are sampled from the Grey matter mask used by the ALE algorithm.

#### 2.3 Algorithm to compute FSN in coordinate-based meta-analyses

Performing an ALE meta-analysis can be time-demanding. In order to reduce the amount of steps necessary to ascertain the FSN it is necessary to define a minimum and maximum value before computing the FSN. There are no formal guidelines for the minimum and maximum FSN, researchers should determine what seems practically relevant for your meta-analysis based on the area of research. A recent study estimated that in fMRI, one out of ten studies remains in the file drawer (Samartsidis, Fox, Laird, Johnson, & Nichols, 2017). Therefore, looking for a FSN between 2k (with k being the number of experiments in a meta-analysis) and 10k seems reasonable. For reference, Rosenthal (1979) used 5k + 10 as a minimum for the FSN. An ALE meta-analysis usually contains between 20 and 30 individual experiments. In general, a FSN of at least 50 seems reasonable.fMRI studies require additional hardware and skills, often making them more expensive than behavioural studies. Therefore it seems reasonable that fMRI studies suffer less from the file drawer problem than behavioural studies. Further simulations are necessary to study the behaviour of the FSN and determine a realistic minimum for fMRI studies. In this paper we aim to introduce the concept of the FSN for fMRI studies. An important remark needs to be made with respect to interpretation of the obtained FSN. For every paper usually several experiments are entered into the meta-analysis, implying that these papers have a stronger influence on the end result than papers that only report 1 experiment. As it is difficult to discriminate between experiments and studies once the list of foci is entered, the FSN is computed on experiment level. The FSN is therefore the number of experiments, and not the number of papers, not contributing to a certain effect that can be added before a cluster is no longer statistically significant. Ideally only one experiment per paper should be added to the meta-analysis. If this is not te case, the researcher should be aware that some papers have more influence on the results and that the FSN is computed on experiment-level, and not on paper-level.

After running an ALE meta-analysis on the original dataset clusters of interest are identified. First for every cluster an ALE meta-analysis is run while a number of noise studies equal to the minimum FSN is added to the original dataset. If the cluster is no longer statistically significant the FSN is smaller than the predefined minimum. If the cluster is still statistically significant after these noise studies are added, a second analysis is run with the maximum amount of noise studies added. If the cluster is still statistically significant the FSN is larger than the maximum, if it is no longer statistically significant the FSN lies somewhere between the predefined minimum and maximum value. To reduce the amount of steps needed to be taken to determine the FSN the number of noise studies added in the next step is the average of the minimum and maximum FSN. Is the cluster still statistically significant, then the minimum is changed to the number of noise studies that was added. If the cluster is no longer statistically significant this number now becomes the maximum. This process is repeated until the minimum and maximum are only one number apart, and the FSN is known. An illustrated example of this procedure can be found in Figure 3.

**Figure 3.**
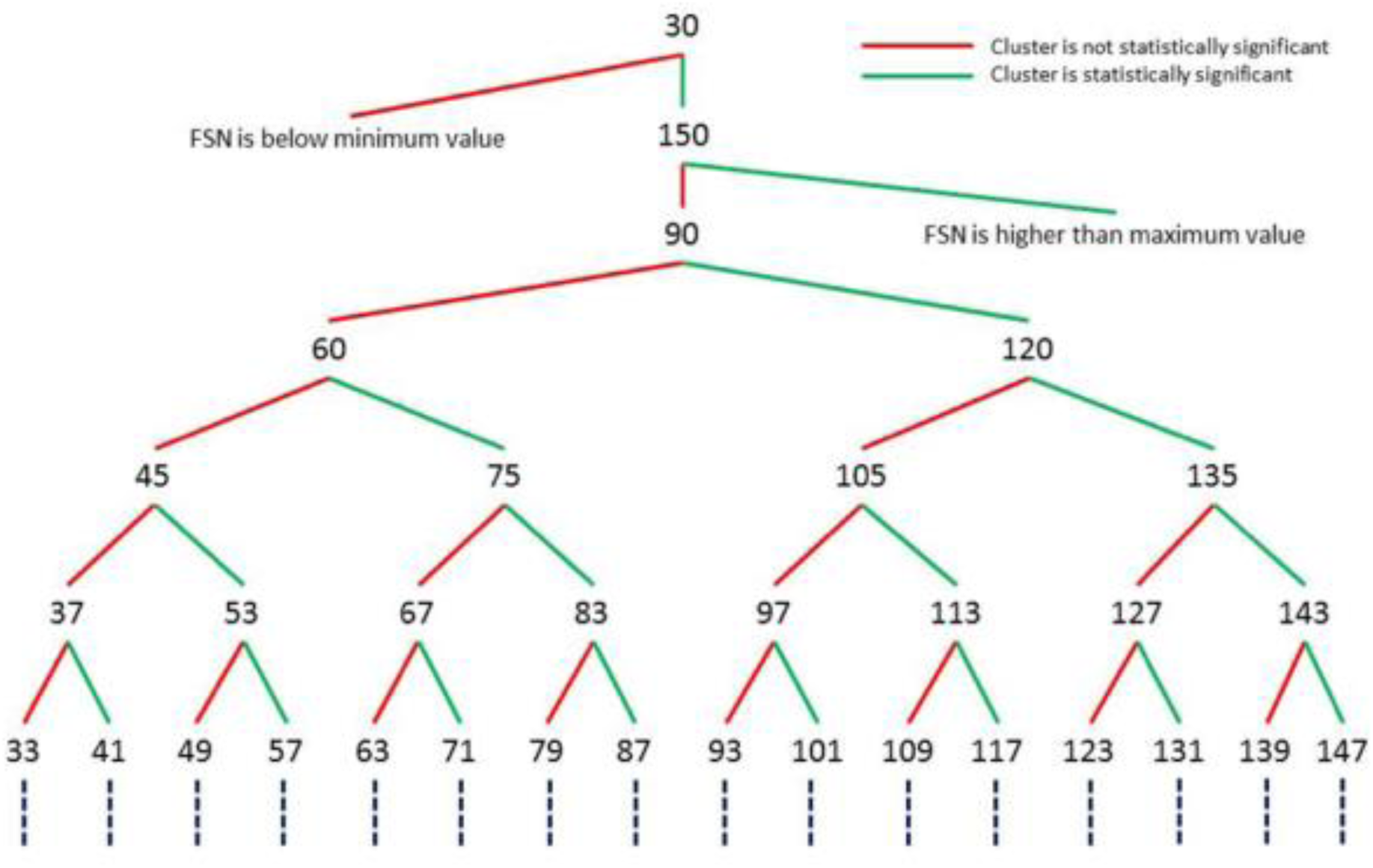
Illustrated example of the algorithm used to compute the FSN of a metaanalysis of 15 experiments (k=15). The predefined minimum is set to 30 (2k) and the predefined maximum is set to 150 (10k). First an ALE meta-analysis is run while a number of noise studies equal to the minimum FSN is added to the original dataset. If the cluster is no longer statistically significant the FSN is smaller than the predefined minimum. If the cluster is still statistically significant after these noise studies are added, a second analysis is run with the maximum amount of noise studies added. If the cluster is still statistically significant the FSN is larger than the maximum, if it is no longer statistically significant the FSN lies somewhere between the predefined minimum and maximum value. To reduce the amount of steps needed to be taken to determine the FSN the number of noise studies added in the next step is the average of the minimum and maximum FSN. Is the cluster still statistically significant, then the minimum is changed to the number of noise studies that was added. If the cluster is no longer statistically significant this number now becomes the maximum. This process is repeated until the minimum and maximum are only one number apart, and the FSN is known.

### 3. Simulations

To study the influence of study parameters, such as sample size, number of peaks per study and thresholding method employed by the ALE algorithm, on the result of the FSN, we simulated meta-analysis sets which consist of lists of peaks, divided into studies. We constructed three versions of 1000 simulated meta-analyses. In each simulation, a meta-analysis of 103 studies is generated, 3 real studies with peaks close to the location of true activation and 100 noise studies that can be added to compute the FSN, that consist of irrelevant random coordinates. The number of real studies is arbitrarily chosen. In a first scenario, every study reports 1 peak, in a second scenario, this is 8 peaks (the average amount of peaks per study entered in BrainMap) and in a third scenario, a random amount of peaks sampled from the BrainMap dataset are reported. The location of these peaks is randomized. To determine the location of the peaks, we divided the brain into 4 quadrants (Figure 4). In quadrant 1 the peaks of the 3 real studies were sampled at a random distance (~N(0,1)) from the location of true activation (MNI: 46, −66, −6). This location was randomly chosen. In version 2 and 3, all peaks were sampled at a short distance from the location of true activation per study. The peaks of the noise studies were randomly selected from a list of voxels inside quadrants 2, 3 and 4 (Figure 4). The simulations are illustrated in Figure 5.

**Figure 4.**
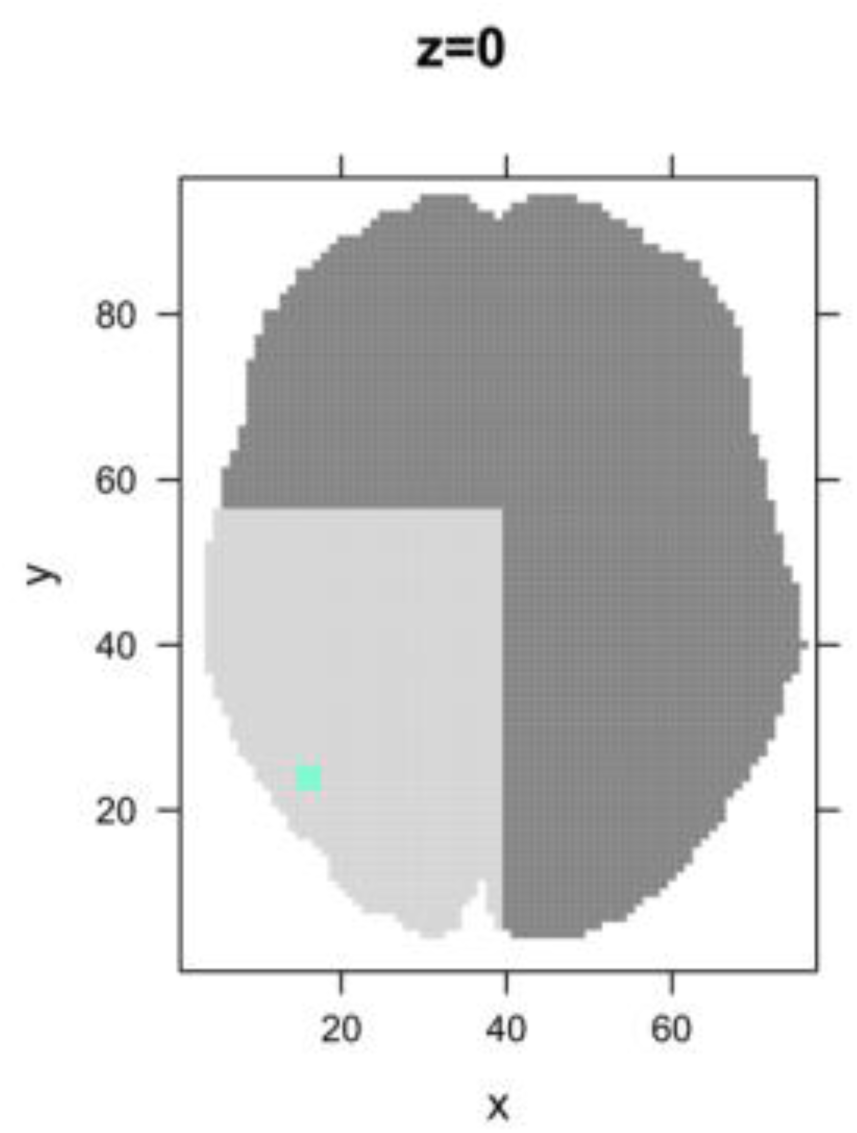
Visualisation of the 4 quadrants the brain is divided into for simulation. The blue dot represents the true location of activation and the peaks of the 3 real studies are sampled around this location. To avoid interference between the real and the noise studies, the peaks of the noise studies are sampled from quadrants 2, 3 and 4 (darker shades of grey).

**Figure 5.**
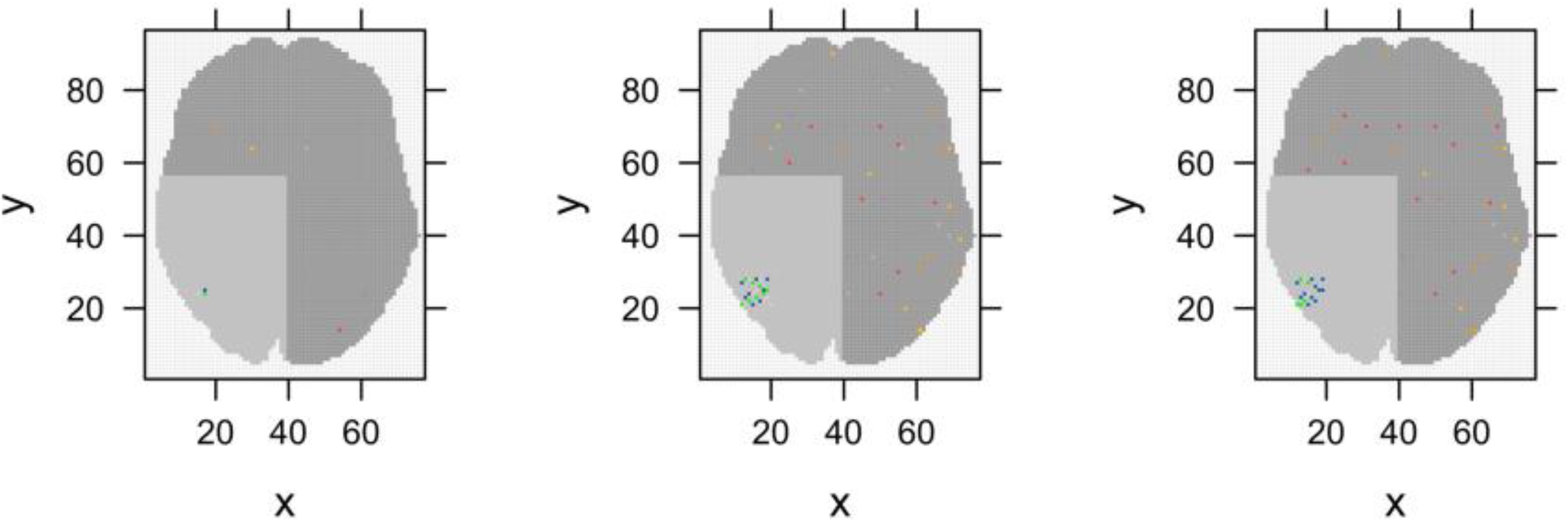
Visualisation of simulation conditions with 3 real studies and 5 noise studies. The visualisation is simplified to 2D slice with z = 0. In the real simulations coordinates are entered into 3D space, resulting in a larger spread. In the first pane we see the scenario with 1 peak per study. The peaks from the 3 real studies lie in quadrant 1, the peaks from the noise studies lie in the other 3 quadrants. The pane in the middle represents the scenario with 8 peaks per study. The real studies each have eight peaks in quadrant 1, resulting in 24 peaks close to eachother. The 5 noise studies each have 8 peaks in the remaining 3 quadrants. Their peaks will never lie in quadrant 1. Due to the random location of peaks some overlap and spurious activation might occur. The influence of this spurious activation is minimal after 1000 simulations. In the right pane the scenario with a random number of peaks is shown. The 3 real studies have 5,11 and 2 peaks respectively. The 5 noise studies have peaks in the remaining quadrants, with 2, 12, 4, 11 and 5 peaks respectively.

To evaluate the effect of sample size, 3 copies of all 1000 sets were created. The only difference between these sets was the average sample size. The sample size of the 3 real studies varied and was randomly sampled (~N(0,1)) around 10 (small sample size), 20 (average sample size) or 30 (large sample size). The sample size of the smallest of the 3 real studies was used as the sample size of the noise studies and remained the same over all 100 noise studies. This was done because small studies use a larger FWHM and therefore have slightly more influence on the end result. This ensured that the noise studies had at least as much weight as all real studies. To compute the FSN, ALE meta-analyses are performed with a varying amount of noise studies until the tipping point of statistical significance is found. Data obtained through simulations was used for validation and the matlab implementation of ALE was used (Eickhoff, personal communication). We applied 3 different methods for thresholding, namely uncorrected (p < 0.001), voxel-wise Family-Wise Error Rate (vFWE, p < 0.05) and a cluster-level FWE thresholding (p < 0.05) with uncorrected cluster-forming threshold (p < 0.001). For the voxel-level thresholding methods no additional minimum cluster size was employed. We analysed 1000 meta-analyses for every thresholding method. 100 permutations within the clustering algorithm were performed for the latter. Datasets and code are available on GitHub (https://github.com/NeuroStat/FailSafeN).

Results are reported in Table 3 and visualised in Figure 6, Figure 7 and Figure 8. A difference in FSN is seen for the choice of thresholding method, the sample size of the individual studies and the amount of peaks. Overall, uncorrected thresholding is most lenient, the FSN was larger than 100 regardless of number of peaks and individual study sample size, and we strongly advise against using uncorrected thresholding for a meta-analysis. The results for voxel-and cluster-level Family-Wise Error Rate differ strongly from the results for uncorrected thresholding, but are comparable to one another, especially in the most ecologically valid scenario with a random number of peaks per study. In the scenario of 1 peak per study and voxel-level Family-Wise Error Rate correction on average 3% of the studies in the meta-analysis stem from the original meta-analysis, the rest are noise studies. However, in the other two cases this is on average between 6% and 9%. In every condition less than 1/10 studies contributes to the activated cluster. A possible explanation for the robustness of the results is that the reported peaks lie close to eachother, which results in a large ALE-value.

**Figure 6.**
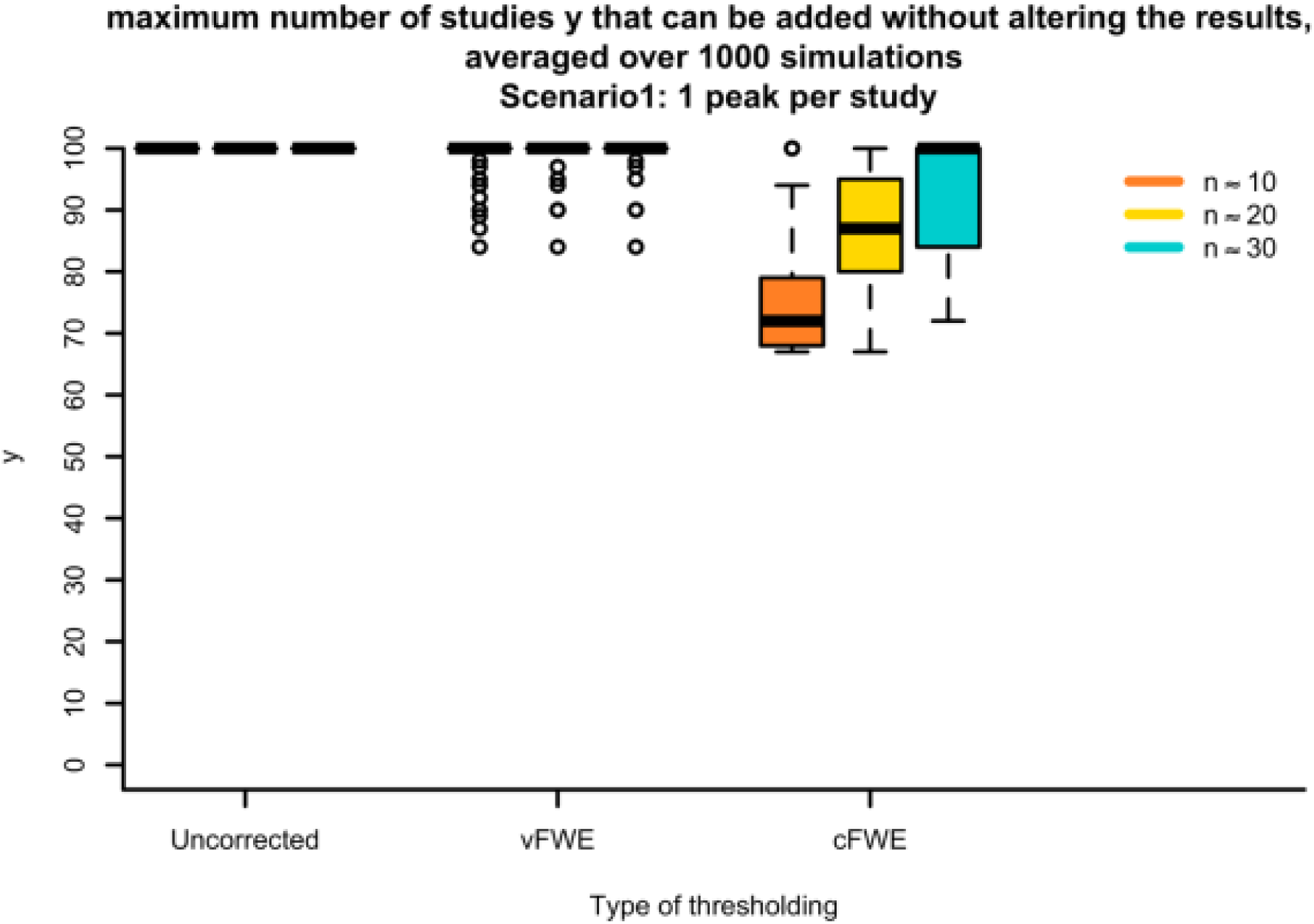
Boxplots of simulations with the FSN in the scenario of one peak per study. On the x-axis the thresholding method and average amount of participants per experiment can be found. On the y-axis the average amount of noise studies that can be added to a meta-analysis of 3 studies with real activation before the target area is no longer statistically significant is plotted.

**Figure 7.**
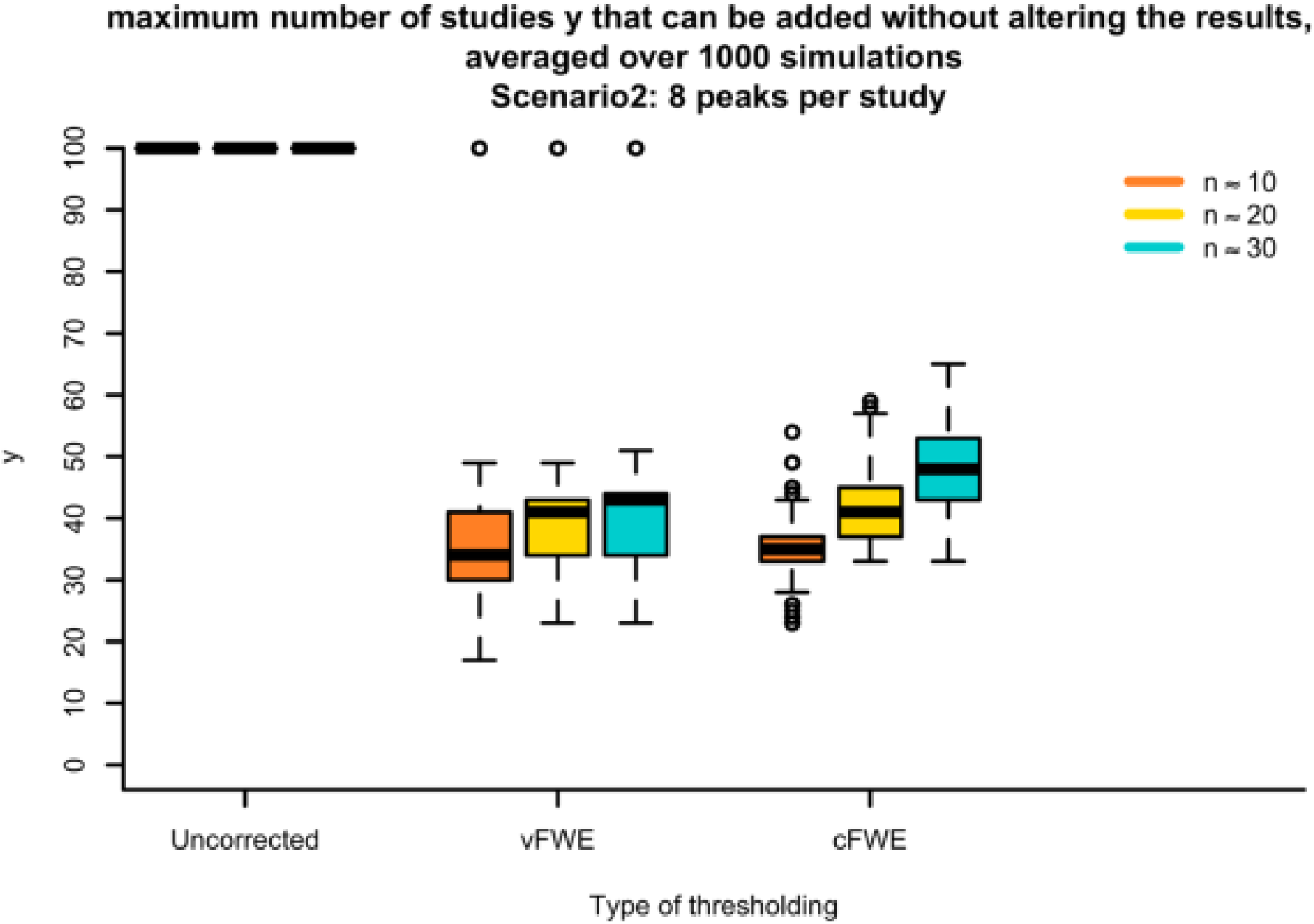
Boxplots of simulations with the FSN in the scenario of eight peaks per study. On the x-axis the thresholding method and average amount of participants per experiment can be found. On the y-axis the average amount of noise studies that can be added to a meta-analysis of 3 studies with real activation before the target area is no longer statistically significant is plotted.

**Figure 8.**
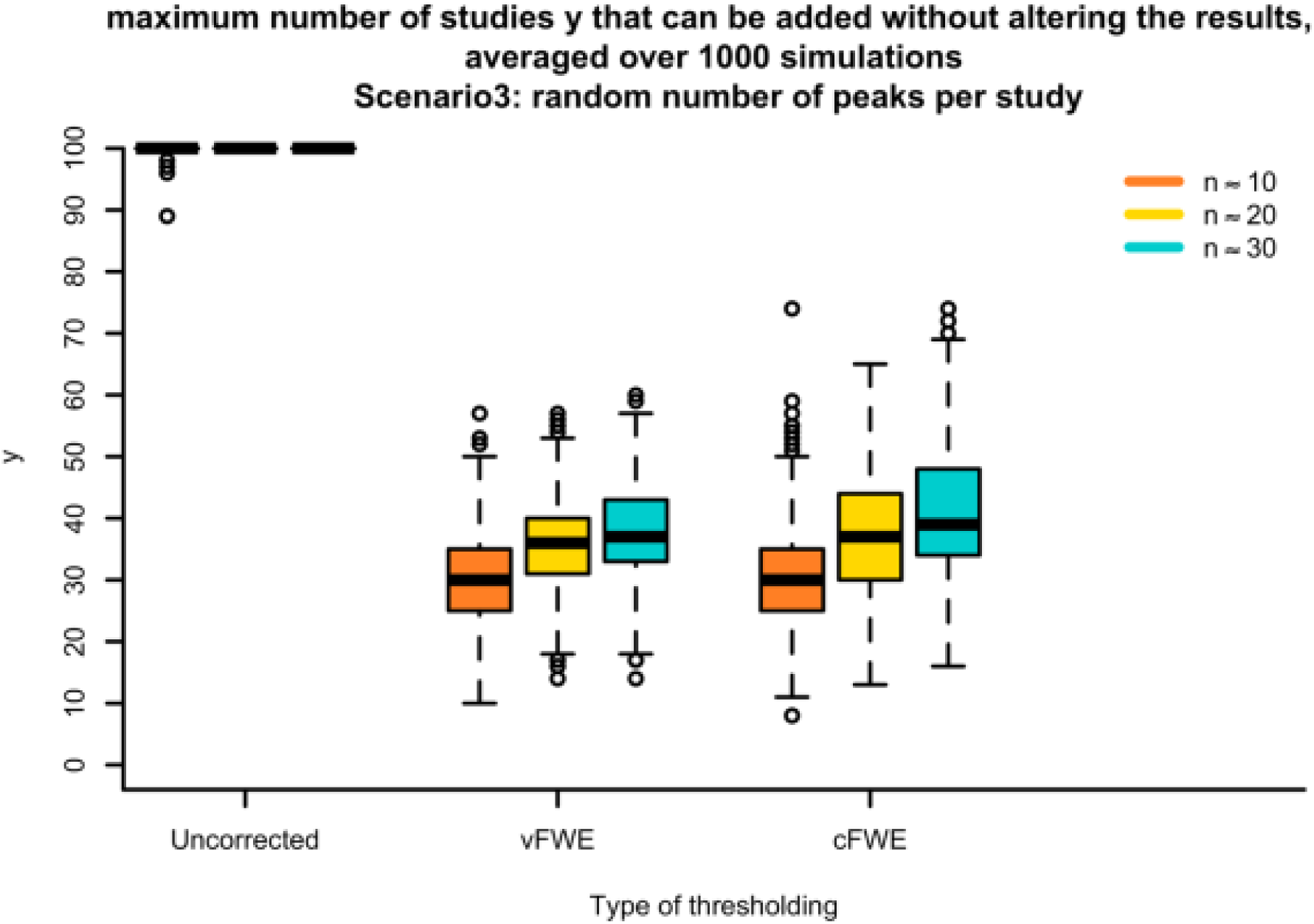
Boxplots of simulations with the FSN in the scenario of a random number of peaks per study. On the x-axis the thresholding method and average amount of participants per experiment can be found. On the y-axis the average amount of noise studies that can be added to a meta-analysis of 3 studies with real activation before the target area is no longer statistically significant is plotted.

A positive effect on robustness can be observed for sample size. It is not possible to view this effect in the case of uncorrected thresholding, but for voxel-and cluster-level Family-Wise Error Rate correction the FSN of meta-analyses with bigger sample sizes is larger. A higher sample size makes the results of a meta-analysis more robust to noise.

The amount of peaks has an important impact on the amount of noise studies that can be added and interacts with the selected thresholding method. Significant effects are mostly driven by a small proportion of studies in the extreme case of 1 peak per study. A ceiling effect can be observed because we did not add more than 100 noise studies. Adding 100 noise studies to a meta-analysis of only 3 studies implicates that less than 3% of the studies in the meta-analysis contribute to a significant effect. A study of the papers present in the BrainMap database shows that for every experiment on average 8 peaks are reported. In this more realistic scenario we see that the FSN drops for every condition and the results for voxel-and cluster-level FWE lie close to one another. The average FSN in the third version, where a random number of peaks sampled from the real distribution of the BrainMap database, does not differ a lot from the average of the second version with 8 peaks per study. There is however a greater variability present in the results, confirming that the results depend heavily on specific parameters of the meta-analysis.

### 4. Example

To demonstrate the principle of the FSN we replicate a previously executed meta-analysis. We selected a meta-analysis on finger tapping (Laird et al. (2008); Eickhoff et al. (2009)) that was used to demonstrate the working of GingerALE from the BrainMap database (Laird et al., 2005; Fox and Lancaster, 2002; Fox et al., 2005). In total 38 papers reporting 73 individual experiments (347 subjects) with a total of 654 activation foci were obtained.

The original meta-analysis consists of 38 papers with k = 73 individual experiments. It is important to note that usually multiple experiments are included per paper. Every experiment is entered into the meta-analysis as a new study. Therefore papers that report multiple experiments have a stronger influence on the results than papers that only report 1 experiment. In the ideal case only one experiment per paper is included. We define a noise study as an experiment, independent of the amount of papers.

Foci from the individual studies were obtained from the BrainMap database. The list of foci was entered into GingerALE 2.3.6 and a meta-analysis was performed. The analysis was thresholded with a cluster-level FWE correction of p < 0.05 (cluster-forming threshold of p < 0.001, uncorrected). After thresholding 7 statistically significant clusters were found, these are reported in Table 1. The location of these clusters is plotted in Figure 9. The clusters varied in size from 27736mm^3^ to 1968mm^3^.

**Table 1.**
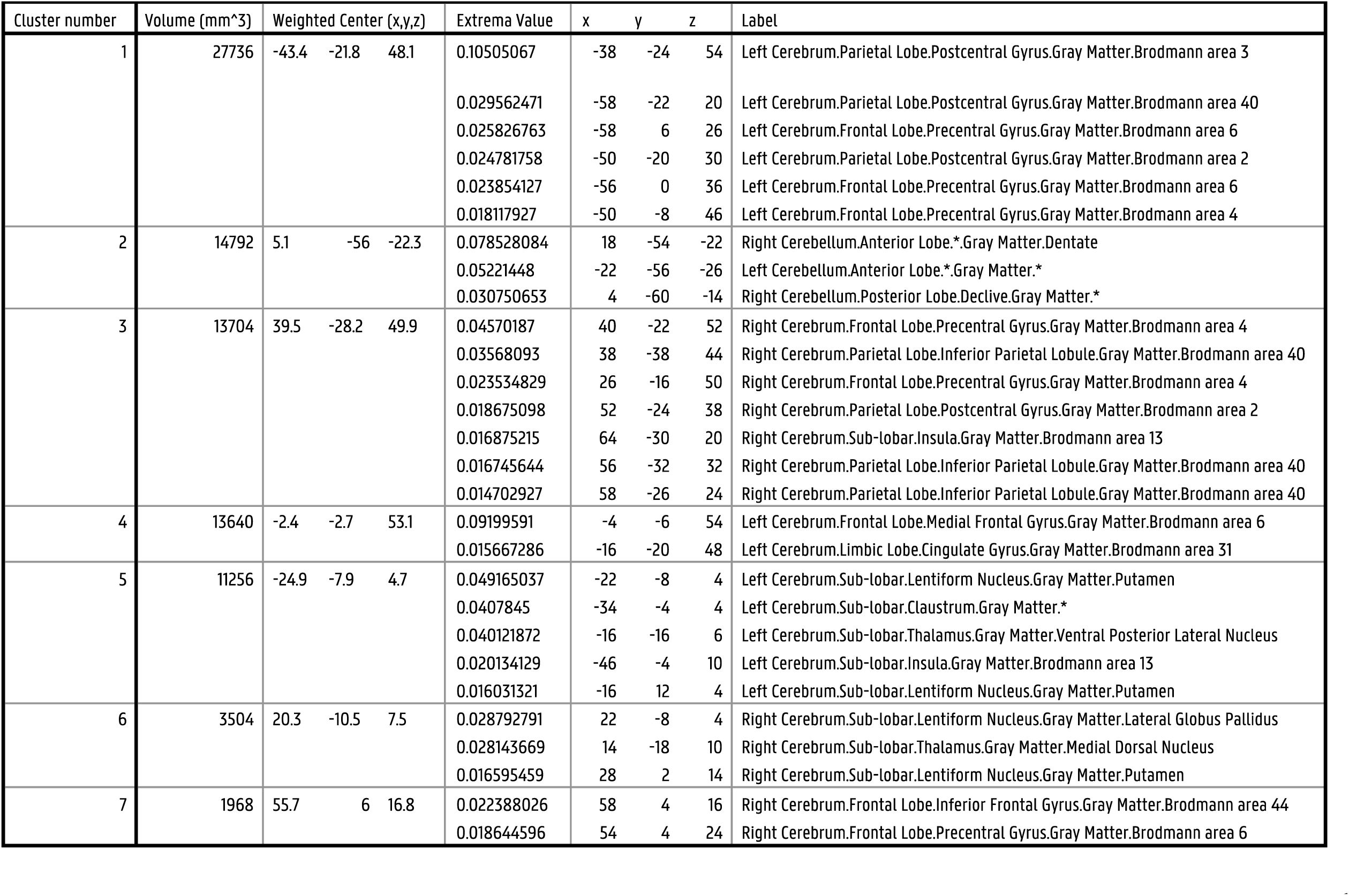

**Figure 9.**
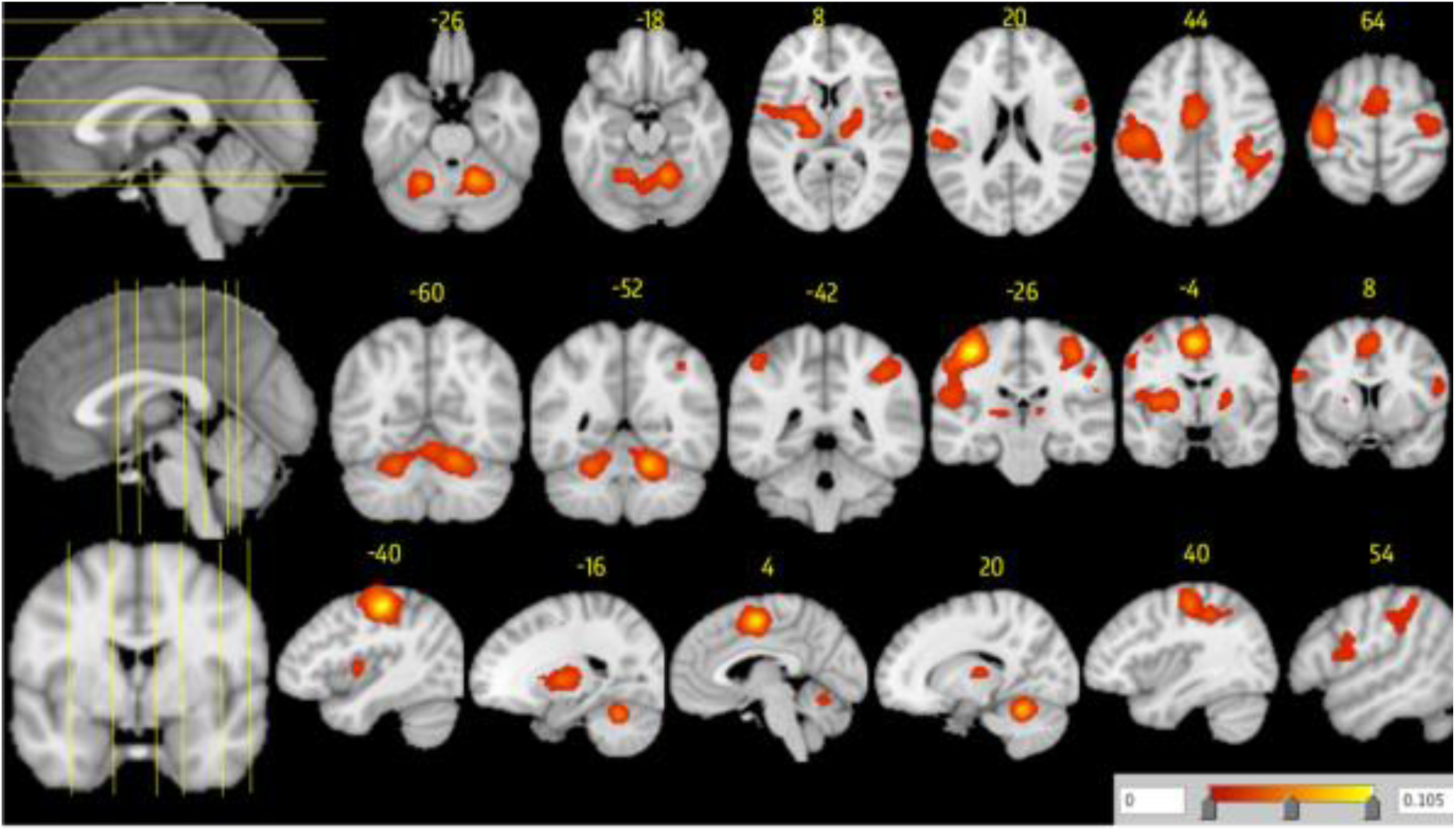
ALE maps were computed using GingerALE 2.3.6 with a cluster-level forming threshold of p < 0.05 (cluster-forming threshold p < 0.001, uncorrected), visualised with Mango.

We continue by computing the FSN for every cluster. We set the minimum of the FSN to 156 (2k + 10) and the maximum to 730 (10k). In this example we first performed an analysis with 156 added noise studies for every cluster. Clusters 6 and 7 were no longer statistically significant if 156 noise studies were added, and have an FSN smaller than the minimum. We hereafter performed an analysis with 730 added noise studies for clusters 1-5. Clusters 1, 2 and 4 remained statistically significant after 730 noise studies were added and have an FSN larger than the maximum. The FSN of clusters 3 and 5 remain unknown. We repeatedly added or removed noise studies as described in the previous section until the FSN was known for these clusters. Cluster 3 was no longer statistically significant if 260 noise studies were added and the FSN of cluster 5 is 409. The FSN for every cluster is displayed in Table 2.

**Table 2.**

| Cluster number | Volume (mm<sup>3</sup>) | Weighted Center (x,y,z) |  |  | Extrema Value | Number of contributing studies | FSN |
| --- | --- | --- | --- | --- | --- | --- | --- |
| 1 | 27736 | -43.4 | -21.8 | 48.1 | 0.10505067 | 59 | > 730 |
| 2 | 14792 | 5.1 | -56 | -22.3 | 0.078528084 | 45 | > 730 |
| 3 | 13704 | 39.5 | -28.2 | 49.9 | 0.04570187 | 42 | 260 |
| 4 | 13640 | -2.4 | -2.7 | 53.1 | 0.09199591 | 53 | > 730 |
| 5 | 11256 | -24.9 | -7.9 | 4.7 | 0.049165037 | 29 | 409 |
| 6 | 3504 | 20.3 | -10.5 | 7.5 | 0.028792791 | 13 | < 156 |
| 7 | 1968 | 55.7 | 6 | 16.8 | 0.022388026 | 10 | < 156 |

**Table 3.**

| Type of thresholding | Number of peaks per study |  |  |  |  |  |  |  |  |
| --- | --- | --- | --- | --- | --- | --- | --- | --- | --- |
|  | 1 peak per study |  |  | 8 peaks per study |  |  | Random number of peaks per study |  |  |
|  | Average sample size |  |  | Average sample size |  |  | Average sample size |  |  |
|  | 10 | 20 | 30 | 10 | 20 | 30 | 10 | 20 | 30 |
| Uncorrected | 100.00 (0.00) | 100.00 (0.00) | 100.00 (0.00) | 100.00 (0.00) | 100.00 (0.00) | 100.00 (0.00) | 99.98 (0.39) | 100.00 (0.00) | 100.00 (0.00) |
| vFWE | 99.75 (1.41) | 99.86 (1.14) | 99.87 (1.18) | 38.48 (18.32) | 40.18 (12.30) | 41.56 (12.45) | 30.05 (7.38) | 35.74 (7.13) | 37.73 (7.39) |
| cFWE (unc) | 75.81 (9.32) | 87.54 (7.74) | 92.59 (9.02) | 34.85 (4.02) | 41.69 (6.33) | 47.06 (6.38) | 30.59 (7.87) | 37.15 (9.17) | 40.84 (10.35) |

We will now look into the results in more detail. We verified which peaks contributed to a cluster and to which experiment these peaks belong. 59 of the 73 experiments in the meta-analysis contributed to cluster 1, explaining why this cluster is very robust to noise and has a FSN larger than the predefined maximum. However, after adding 730 noise studies only 7 of the studies in the meta-analysis contribute to this cluster. If the noise studies are added the cluster is still statistically significant even if it is driven by a small number of studies. Cluster 2 and 4 had 45 and 53 contributing experiments respectively and also had an FSN larger than the maximum. But after adding the maximum number of noise studies less than 6 % of the studies in the meta-analysis contribute to these clusters. This is not the case for cluster 3. Even though 42 out of 73 experiments contributed to this cluster, it has a FSN of 260, much smaller than the maximum. If the maximum number of null studies is added before the cluster is no longer statistically significant, still 13 % of the studies in the meta-analysis contribute to this cluster. A similar result is seen for cluster 5. With 29 contributing studies in the meta-analysis it has a FSN of 409. However, if the 409 noise studies are added to the meta-analysis, only 6% of the studies contribute to the activated cluster. Cluster 6 and 7 had a FSN smaller than the minimum of 156, with 13 and 10 contributing studies respectively. We cannot be certain about the percentage of contributing experiments, but it lies between 5% and 18% for cluster 6 and between 4% and 14% for cluster 7.

We interpreted the results in terms of experiments, but for most fMRI studies a paper consists of several experiments. A lot of variability is present in the amount of experiments per paper. In this meta-analysis there are on average 2 experiments per paper. In terms of papers we can estimate that clusters 1, 2, 3 and 4 are resistant to noise from 365 unavailable papers, cluster 5 to 205 unavailable papers and clusters 6 and 7 to less than 78 unavailable papers.

### 5. Discussion

In this study we focus on a specific form of publication bias, the file-drawer problem. Publication bias occurs when the results of published and unpublished studies differ significantly. In the case of the file-drawer problem studies that do not show statistically significant results or show results that do not match the researchers hypothesis fail to get published. In a most extreme case all studies that remain unpublished are noise studies. We proposed a method that implements the principles of the FSN for the meta-analysis of fMRI studies. Through simulations we show the influence of different parameters on the FSN and through an example we show how we can compute the amount of noise studies that can be added to a meta-analysis before a cluster of interest is no longer statistically significant.

The results of the simulations show that thresholding method, individual study sample size and number of peaks have a large influence on the resulting FSN. In accordance with general guidelines Eickhoff et al. (2016) it is shown that uncorrected thresholding is a very lenient threshold and should not be employed. Also in accordance with this paper we see that voxel-and cluster-level Family-Wise Error correction is better at reducing the excessive contribution of individual studies on the final results than uncorrected thresholding, while still maintaining high power.

The results also show the importance of large sample sizes for the robustness of the results. fMRI studies tend to employ small sample sizes, mainly due to the high costs of executing a study, but this has two main disadvantages. First the power will subsequently drop significantly, making it more difficult to detect an effect. As typically only peak locations are reported this increases the chance of consistent small effects remaining undetected, even if a meta-analysis is executed. Second, small studies are prone to employ a more lenient threshold to compensate for low power. This leads to an increase in false positives and more noise in the meta-analysis.

We also note that the number of peaks has a significant influence on the FSN. Depending on i.e. the software used for the fMRI data-analysis and the thresholding method, some papers consistently report more peaks than others, and therefore have a bigger influence on the results of the meta-analysis. While some studies report 1 activated peak for a relatively large cluster, other studies might report 3 peaks inside a relatively small cluster. The ALE algorithm compensates this by taking the maximum of overlapping kernels within the same study (in the MA-map), but it is impossible to completely neutralize this reporting bias. Moreover, reporting a small amount of peaks is worse for the meta-analysis results than reporting a large amount of peaks.

These results lead to the conclusion that the resulting FSN depends heavily on the individual study parameters. Some other simulation parameters might have influenced the results. One possible parameter is the number of contributing studies. We did not vary the number of real studies in the meta-analysis. If more studies contribute to a statistically significant cluster, the cluster will be more robust to noise. A second possible parameter is the spread and distance of the contributing peaks. Peaks that lie close to each other will result in smaller clusters with higher ALE-values, therefore being more robust to noise. A third and final simulation attribute is the lack of noise peaks in the quadrant of the target cluster. All noise peaks are generated at a location sufficiently far from the cluster of interest, avoiding interference. However, real noise is expected to be spread out throughout the whole brain. We did not verify the influence of noise peaks overlapping with our cluster of interest, but Eickhoff et al. (2016) have shown that this influence is minimal.

To deal with the possible influence of meta-analysis parameters we developed a tool to generate noise studies that are adapted to a researcher’s specific meta-analysis. By re-using the number of foci and sample sizes from the original studies the set of noise studies becomes more ecologically valid. In this tool noise is also spread throughout the brain.

But why should a researcher compute the FSN after conducting a coordinate-based meta-analysis? After entering a list of foci, and losing more than 99% of information obtained by the individual studies, a map with clusters that survived thresholding is received as output. The location of these clusters can easily be retrieved, but interpreting their robustness has proven to be more difficult. Researchers do not always verify the height of the ALE-values in these clusters and the number of studies and foci that contributed to them. The FSN is a useful tool that facilitates determining the robustness of a statistically significant cluster. Researchers can now, after they computed a meta-analysis, download an R program from GitHub (https://github.com/NeuroStat/GenerateNull). In this program they only need to enter the same list of foci they originally used in the meta-analysis. Automatically a list of 10k (with k the number of experiments in the original meta-analysis) noise studies is generated. This number can be adapted if necessary. These noise studies are outputted in the format necessary for the ALE algorithm and can easily be added to the original list of foci. By adding a predefined minimum and maximum number of noise studies and subsequently adding or removing noise studies the FSN can be computed. The algorithm for reducing computational time most efficiently is extensively described in the documentation that can be found on GitHub. When the FSN is found for every cluster the researcher can interpret their robustness to noise and their possible sensitivity to a publication bias.

One might wonder whether the FSN is the optimal tool in this situation. Even though it was quite popular when it was first developed by Rosenthal 1979, because of its ease to use and interpret, it is rarely employed in classic meta-analyses today. The FSN was mainly criticized for the variety in methods and definition of null studies, resulting in a large range of possible results (Rothstein et al., 2006; Scargle, 2000; Schonemann and Scargle, 2008). This critique does not apply in the meta-analysis of fMRI studies. As it is not possible to add a null study where not a single voxel survived statistical thresholding, a null study is defined as a study with random activation, and called a noise study. From a theoretical point of view this makes sense. As more than 100.000 statistical tests are performed, some spurious activation can be expected. This is reflected in the random peaks that are simulated. Specifically for the ALE-algorithm this is a logical rationale because it looks for convergence in activation. Randomly sampled peaks by definition show no convergence in activation and are therefore an adequate representation of a noise study. The FSN also received critique for its focus on statistical significance. While we strongly recommend the use of effect sizes, it is not possible to employ effect sizes in this situation as only the location of activation is given. We merely wish to provide tools for researchers that facilitate the interpretation of meta-analytical results. The over-abundant focus on p-values is an issue inherently linked to science. By providing the FSN as a tool we want to encourage researchers to look for something else than the statistical significance of a region of interest, such as size and location of the cluster, number of contributing studies and foci and robustness to noise. As the results of an ALE meta-analysis depend solely on the location of reported activation foci and the sample size of individual studies, a researcher should be aware of the number of studies, and the sample sizes of these studies, that contributed to an activation cluster. This data is listed in the results of an ALE meta-analysis. With the aid of the FSN the researcher can determine the robustness to noise of a cluster of interest, and interpret these results in the light of number of contributing foci and studies.

We would also like to note that combining entire statistical parametric maps (SPM) through a random-effects meta-analysis is preferred over a meta-analysis where merely peaks are entered. However, for the majority of fMRI studies these SPM’s are not readily available. We therefore need coordinate-based methods such as the ALE and MKDA algorithm and our study provides means to assess publication bias and verify robustness of clusters resulting from a coordinate-based meta-analysis.

Publication bias is an issue that has been discussed in science since the 18th century (Ferriar, 1792; Sterling, 1959; Hall, 1965) where researchers note that journals are mainly filled with papers that report statistically significant results. Possible explanations include that journals are less likely to accept papers with null results and that researchers are hesitant to submit a paper that is not in line with their research hypothesis. The FSN was originally developed to assess the impact of the file-drawer problem, where studies that fail to show statistically significant results remain in the file drawer. It is difficult to assess the severity of this issue, we can never be sure of the amount of (well executed) studies that never got published, and of the results they report. We can only hope that the amount of unpublished studies is small and that their results are similar to those of published studies. If this is not the case, the FSN gives an estimation of the number of unpublished studies with different results that would be needed to change the conclusions of a meta-analysis on a given effect.

While we focussed on the file-drawer problem, other forms of publication bias exist as well. The FSN assesses the impact of a between-study publication bias, but the impact of a within-study publication bias cannot be ignored. While large studies have more power to detect true effects, small studies tend to apply more lenient thresholding to compensate a lack of power. Furthermore fMRI studies suffer from small sample sizes, mainly because they are very costly and time-demanding. This may lead to a huge increase of false positives. Researchers should therefore verify whether an activated cluster is mainly driven by large or small studies and interpret the results.

Even though researchers should still strive towards making the statistical parametric map of their study available, the ALE algorithm offers a considerable alternative in the meantime. It is important to develop tools to aid researchers with comprehending the meaning of the results of their meta-analysis. The FSN offers researchers an opportunity to verify the robustness of the statistically significant clusters and the conclusions they deduce from these clusters based on parameters stemming from the original meta-analysis.

## Conclusion

Introducing an adaptation of the Fail Safe N (FSN) in the meta-analysis of fMRI studies greatly aids researchers in assessing robustness of fMRI meta-analysis results. This is the first tool that allows assessing cluster robustness to noise and the cluster’s sensitivity to publication bias. For every cluster resulting from the fMRI meta-analysis the FSN can be computed, showing the number of noise studies that can be added before the cluster is no longer statistically significant. We provide a tool that generates noise studies that are based on the number of foci and sample sizes of the studies that are originally entered into the meta-analysis. This tool is accompanied by a step-by-step guide on how to compute the FSN, facilitating it’s computation. The process is further illustrated in a comprehensive example provided in this manuscript.

We additionally show the influence of different study parameters on the FSN through extensive simulations. The simulations confirm that robustness increases with sample size and that number of foci has a strong influence on the FSN. By showing the influence of individual study parameters such as number of foci and sample size on the FSN, we mark the importance of generating noise studies that are similar to the original studies. The simulations also confirm that uncorrected thresholding should not be employed with the ALE algorithm, which is in line with the conclusion from a massive empirical simulation study (Eickhoff et al., 2016). In conclusion, the adaptation of the FSN for fMRI meta-analysis provides a measure for robustness of resulting clusters and their sensitivity to publication bias which provides an added value to the interpretation of brain regions selected as statistically significant by the ALE algorithm.

## Acknowledgments

FA, RS and BM would like to acknowledge the Research Foundation Flanders (FWO) for financial support (Grant G.0149.14).

SBE was supported by the National Institute of Mental Health (R01-MH074457), the Helmholtz Portfolio Theme “Supercomputing and Modeling for the Human Brain” and the European Union’s Horizon 2020 Research and Innovation Programme under Grant Agreement No. 7202070 (HBP SGA1).

